# Flanker task parameters are related to the strength of association between the ERN and anxiety: a meta-analysis

**DOI:** 10.1101/2024.08.27.609944

**Authors:** George A. Buzzell, Yanbin Niu, Emily Machado, Renata Dickinson, Jason S. Moser, Santiago Morales, Sonya V. Troller-Renfree

## Abstract

The error-related negativity (ERN)—an index of error monitoring—is associated with anxiety symptomatology. Although recent work suggests associations between the ERN and anxiety are relatively modest, little attention has been paid to how variation in task parameters may influence the strength of ERN-anxiety associations. To close this gap, the current meta-analysis assesses the possible influence of task parameter variation in the Flanker task—the most commonly used task to elicit the ERN—on observed ERN-anxiety associations. Here, we leveraged an existing open database of published/unpublished ERN-anxiety effect sizes, supplementing this database by further coding for variation in stimulus type (letter vs. arrow), response type (one-handed vs. two-handed), and block-level feedback (with vs. without). We then performed meta-regression analyses to assess whether variation in these Flanker task parameters moderated the effect size of ERN-anxiety associations. No evidence for an effect of stimulus type was identified. However, both response type and block-level feedback significantly moderated the magnitude of ERN-anxiety associations. Specifically, studies employing either a two-handed (vs. one-handed) task, or those with (vs. without) block-level feedback exhibited more than a two-fold increase in the estimated ERN-anxiety effect size. Thus, accounting for common variation in task parameters may at least partially explain apparent inconsistencies in the literature regarding the magnitude of ERN-anxiety associations. At a practical level, these data can inform the design of studies seeking to maximize ERN-anxiety associations. At a theoretical level, the results also inform testable hypotheses regarding the exact nature of the association between the ERN and anxiety.

---

The error-related negativity (ERN) is a response-locked ERP characterized by a frontocentral negative voltage deflection occurring approximately 0-100 ms after committing an error (Gehring et al., 2012). Several meta-analyses suggest that a larger (i.e., more negative) ERN is robustly associated with anxiety symptomatology (Cavanagh & Shackman, 2015; Moser et al., 2013; Pasion & Barbosa, 2019; Riesel, 2019; Saunders & Inzlicht, 2020). However, one recent meta-analysis found that, after accounting for possible publication bias, the magnitude of the ERN-anxiety association may be relatively modest (Saunders & Inzlicht, 2020). Yet, prior work has been limited in considering how task variation across studies might impact meta-analytic estimates of ERN-anxiety associations. Prior meta-analyses report a fair degree of homogeneity in the kind of tasks employed for studying ERN-anxiety associations: most studies employ conflict-related tasks, with the Flanker task being most the most common (e.g., Moser et al., 2013; Saunders & Inzlicht, 2020). Nevertheless, several common variations of the Flanker task (Eriksen & Eriksen, 1974) have been employed across studies. Despite recent calls to directly assess the impact of variation in task parameters on ERN-anxiety associations (Gloe & Louis, 2021), few studies have done so. Moreover, there is no meta-analytic data speaking to whether variation in stimulus, response, or feedback parameters influence the strength of estimated ERN-anxiety associations.

The goal of the current meta-analysis was to assess the possible influence of variation in Flanker task parameters on observed ERN-anxiety associations. This goal is important for several reasons. First, by accounting for variation in task design, it may be possible to at least partially account for apparent inconsistencies in the literature regarding the strength of ERN-anxiety associations. Second, and at a practical level, identifying which task parameters maximize ERN-anxiety associations could inform the design of future studies seeking to examine this effect. Third, while the current meta-analysis cannot directly test competing theories on the nature of ERN-anxiety associations, results could inform the development of testable hypotheses regarding how and why the ERN is associated with anxiety. Ultimately, this work can advance the theoretical understanding and the clinical relevance of the ERN to anxiety problems.

Our rationale for which task parameters to examine was based on practical limitations and prior theory of the established literature. Given that most studies investigating ERN-anxiety associations used a Flanker task (79% of studies reported in the recent meta-analysis by Saunders & Inzlicht, 2020), we focused our analyses on studies that used this task to hold the larger task structure constant while assessing whether specific task parameters influenced ERN-anxiety associations. Within these constraints and further informed by prior theory, we identified three parameters of interest that would be feasible to analyze in the current meta-analysis: stimulus type (arrow vs. letter), response type (one-vs. two-handed), and block-level feedback (with vs. without). Below, we briefly review relevant theory supporting the possibility that these parameters might impact ERN-anxiety associations.

Variation in stimulus type (e.g., letters vs. arrows) could impact ERN-anxiety associations for at least two reasons. One possibility is that differences in the strength of stimulus-response mappings for letter vs. arrow stimuli could impact ERN-anxiety associations. For instance, for letter stimuli, stimulus-response mappings are arbitrary and there is no preexisting relationship between a given letter stimulus and a particular response option (button press). In contrast, for arrow stimuli, there is a preexisting semantic-spatial compatibility (e.g., pressing the right response button for a right-facing arrow; Kornblum et al., 1990). Thus, performing an arrow Flanker task might place less demands on working memory and allow for more efficient/automatic response preparation (Kornblum et al., 1990; Wilhelm & Oberauer, 2006). These differences could impact error monitoring, in turn influencing ERN-anxiety associations. An alternative possibility is that linguistically oriented stimuli (letters) may be more likely to engage language processing (Jacobs & Grainger, 1991; Petit et al., 2006) and prime verbal rumination, ultimately increasing ERN-anxiety associations. One prominent theory, the Compensatory Error-Monitoring Hypothesis (CEMH) of Moser and colleagues (Lin et al., 2015; Moser et al., 2013), proposes that ERN-anxiety associations are driven by the presence of verbal rumination. Thus, it is possible that the use of more linguistically oriented stimuli (letters) could elicit stronger ERN-anxiety associations.

Variation in response type (whether participants respond to stimuli using one hand or two) could lead to differences in ERN-anxiety associations for multiple reasons. As previously mentioned, the CEMH of Moser and colleagues (Lin et al., 2015; Moser et al., 2013) proposes that ERN-anxiety associations are driven by the effects of verbal rumination. Specifically, this theory proposed that, when performing a two-handed task, verbal rumination primes left motor cortex (Engels et al., 2007; Oathes et al., 2008), in turn increasing the relative prepotency of right-vs. left-hand responses (Lin et al., 2015). As a result, when high anxious individuals perform a two-handed task, errors made with the right hand are more prepotent and require greater inhibition (eliciting a larger ERN; Hochman et al., 2014; Lin et al., 2015). Given that the CEMH theory is based on the notion that *relative* asymmetries in right/left responses drive this effect (Hochman et al., 2014), the theory would predict that such effects would only be present in two-handed, not one-handed tasks, even if all responses were made by the right hand (Lin et al., 2015). This is because only in a two-handed task can left-lateralized verbal rumination lead to a *relative* increase (difference) in the prepotency between right (vs. left) hand responses. Thus, the CEMH theory (Lin et al., 2015; Moser et al., 2013) would predict stronger ERN-anxiety associations for two-handed vs. one-handed tasks.

A separate possible reason for why response type could impact ERN-anxiety associations draws on a longstanding debate over whether the ERN arises from conflict—as described by conflict monitoring theory (Botvinick et al., 2001; Yeung et al., 2004)—or as the result of a mismatch detection process, as described by comparator/mismatch theories of the ERN (Bernstein et al., 1995; Coles et al., 2001; Falkenstein et al., 1996). At least some interpretations of conflict monitoring theory predict greater conflict—and a larger ERN—for more similar responses (e.g., fingers on the same hand), whereas comparator/mismatch theories predict more salient error detection—and a larger ERN—for more dissimilar responses (e.g., fingers on different hands; Bernstein et al., 1995; Falkenstein et al., 1996; Gehring & Fencsik, 2001). ERN variation may arise from both conflict monitoring and comparator/mismatch detection (Arbel & Donchin, 2011; Bernstein et al., 1995; Falkenstein et al., 1996; Gehring & Fencsik, 2001). Yet, to the extent that anxiety is more strongly associated with ERN variation arising from one process vs. the other, differences in ERN-anxiety associations could emerge as a function of response type.

Finally, a third type of variability that may influence the magnitude of the ERN-anxiety associations is whether block-level feedback is presented to participants. A common method of ensuring that participants make enough errors for ERN analyses is to present block-level feedback contingent on performance. That is, consistent with the recommendations of Gehring and colleagues (2012), researchers often present feedback instructing participants to “go faster” if too few errors are made, and present feedback to “be more accurate” when too many errors are committed. However, the presence of such feedback could also influence ERN-anxiety associations for more than one reason. One possibility is that the presence of any form of feedback could serve as a quasi-social manipulation insofar as it reminds participants that their performance is being monitored/evaluated by researchers (Blair et al., 2008; Hajcak et al., 2005). Given that ERN-anxiety associations are strengthened within social contexts, at least for social anxiety symptoms (Barker et al., 2015; Niu et al., 2023), block-level feedback might also serve to increase ERN-anxiety associations. An alternative possibility is that anxious and non-anxious individuals differentially modulate error monitoring levels in response to the speed vs. accuracy instructions inherent in block-level feedback to “Go faster” vs. “Be more accurate.” Indeed, Riesel and colleagues (2019) found that the ERN of individuals with Obsessive-Compulsive Disorder (OCD) vs. healthy individuals was differentially influenced by speed vs. accuracy instructions. When provided with instructions to respond faster (vs. instructions to respond accurately), healthy individuals exhibited a substantial reduction in their ERN (Riesel et al., 2019). In direct contrast, individuals with OCD showed a more modest reduction in their ERN under speed instructions (Riesel et al., 2019). Thus, non-anxious individuals may be more able to flexibly modulate their error monitoring in accord with task instructions, yielding a reduced ERN within the context of speed (vs. accuracy) feedback. In contrast, anxious individuals may exhibit less variability in their error monitoring levels, rigidly deploying heightened error monitoring regardless of task/feedback instructions. Note that such predictions are also consistent with Attentional Control Theory (Eysenck et al., 2023), which proposes that anxious individuals deploy heightened error monitoring as a compensatory strategy to maintain performance and avoid errors, regardless of task instructions (Eysenck et al., 2023; Moser et al., 2013). Taken together, and given that studies of the ERN commonly employ block-level feedback that includes performance-contingent instructions to “Go faster”, the presence of such feedback may also increase ERN-anxiety associations (for a similar argument, also see: Gloe & Louis, 2021).

In the current study, we performed meta-regression analyses to assess whether variation in stimulus type, response type, or block-level feedback influence the magnitude of ERN-anxiety associations amongst Flanker task studies. To this end, we leveraged an existing open data set of published/unpublished ERN-anxiety effect sizes created by Saunders and Inzlicht (2020). For each Flanker task study listed in the Saunders and Inzlicht (2020) data set, we additionally coded each study in terms of stimulus type, response type, and block-level feedback presence, by referring to the original research publications and/or contacting authors via email. As reviewed in two prior meta-analyses (Moser et al., 2013; Saunders & Inzlicht, 2020), anxiety reflects a multidimensional construct that includes a cognitive component of worry/anxious apprehension, as well as a visceral/autonomic component of anxious arousal/fear among other components. In the current meta-analysis, to maximize the number of included studies, our primary analyses all focused on associations between the ERN and anxiety, broadly defined (i.e., without distinguishing between worry/anxious apprehension and other dimensions of anxiety). Nonetheless, to allow for comparison with prior work, we also performed an exploratory set of analyses restricted to studies employing anxiety measures that mapped onto the anxious apprehension/worry component of anxiety (see supplement). Although our decision of which parameters to analyze was motivated by prior theory, the primary goal of this study was to establish—at the meta-analytic level—whether specific task parameters (stimulus type, response type, block-level feedback) influence the strength of ERN-anxiety associations. Given that our analyses were not intended to provide direct tests of prior theory, we did not have strongly motivated a priori hypotheses as to the direction of observed effects. Nonetheless, in the discussion section we expand on both the practical and potential theoretical implications of the observed results.

## Methods

### Overview

The initial list of studies and all effect sizes included in the current meta-analysis were sourced from an open data set of published/unpublished ERN-anxiety effect sizes created by Saunders and Inzlicht (2020): (https://osf.io/r7dvc/). The aim of the current study was to re-examine these effect sizes in conjunction with new data that our team extracted from the original studies, testing for whether specific task parameters (stimulus type, response type, block-level feedback) moderate the ERN-anxiety associations among Flanker task studies.

### Power analyses

Power analysis performed via the *metapower* package in R (Griffin, 2021) indicated that approximately 26 studies would be needed to achieve >80% power to detect small-medium effect size differences (r = .2) as a function of categorical moderators within a fixed-effects meta-regression framework. This power analysis was informed by the prior meta-analysis by Saunders and Inzlicht (2020), such that we assumed a mean effect size of r = -.19, I^2^ = 52.5%, and a mean study sample size of 66.

### Study selection

#### Prior meta-analysis performed by Saunders and Inzlicht (2020)

The current meta-analysis was performed on a subset of studies taken from the Saunders and Inzlicht (2020) database. Here, we provide a brief overview of the search and selection criteria applied by Saunders and Inzlicht (2020); for complete details, readers should refer to the original paper. Inclusion/exclusion criteria for the Saunders and Inzlicht (2020) dataset were designed to be consistent with two prior meta-analyses (Cavanagh & Shackman, 2015; Moser et al., 2013). To this end, studies were included if they contained a measure of the ERN and a measure of anxiety, broadly defined, to include “studies where groups were defined based on clinical diagnosis (e.g., generalized anxiety disorder, social anxiety disorder, obsessive-compulsive disorder, post-traumatic stress disorder), anxiety scales (e.g., Penn State Worry Questionnaire; Yale-Brown Obsessive Compulsive Scale; Anxiety Sensitivity Index), and a range of closely related individual differences (e.g., Big-5 Neuroticism, Behavioral Inhibition System Scale)” (Saunders & Inzlicht, 2020, p. 89). Additionally, studies were only included if they employed a “conflict paradigm (e.g., Stroop, Flanker, Go/Nogo) with no motivational manipulation (e.g., monetary incentive)” (Saunders & Inzlicht, 2020, p. 89) to be consistent with the meta-analysis conducted by Moser and colleagues (2013). Employing these inclusion criteria, Saunders and Inzlicht (2020) identified published and unpublished studies for inclusion based on: 1) the two prior meta-analyses (Cavanagh & Shackman, 2015; Moser et al., 2013); 2) an open call for published and unpublished studies communicated via email, social media, and OSF; 3) a web-search and reading of systematic reviews; 4) a PubMed search constrained from January 2012 to June 2018. Following these procedures, the final dataset created by Saunders and Inzlicht (2020) was comprised of 58 studies.

#### Study selection criteria for current meta-analysis

In addition to the primary selection criteria imposed by Saunders and Inzlicht (2020), we further restricted study selection to only include studies from the Saunders and Inzlicht (2020) database that used a Flanker task (Eriksen & Eriksen, 1974)—ensuring all studies employed the same overall task structure. Holding task structure constant across all our meta-regression analyses was done to avoid confounding overall task structure with the potential moderating role of each parameter of interest (stimulus type, response type, block-level feedback). The Flanker task was chosen because, 1) a substantial majority (79%) of studies reported in the Saunders and Inzlicht (2020) dataset employed a Flanker task, and 2) all parameters of interest can be coded for this task. To this end, we coded studies in terms of whether they employed a Flanker task to elicit an ERN. Following the exclusion of studies that did not employ a Flanker task structure, 46 studies remained for further analysis (see Table 1). For all published studies, we had two independent coders extract and reconcile these task codes from the original articles. For unpublished studies, task codes were derived from information present within the Saunders and Inzlicht (2020) database. Of the 46 studies coded, we further coded whether the task structure was “modified” to include additional stimuli or response demands that are not typical of most Flanker task studies (see supplement for further details). A subset of 5 studies were identified as being a “modified” version of the standard Flanker task structure. These modified Flanker task studies were included in all primary analyses reported in the main text; however, we additionally re-ran all meta-regression analyses testing for moderation after removing the modified Flanker task studies and obtained similar results (see supplement).

**Table 1.** Studies included in the meta-analysis; adapted from Saunders and Inzlicht (2020), Moser and colleagues (2013), as well as Cavanagh and Shackman (2015).

| First author (year) | Population | Anxiety Measure | Task | Stimulus | Response | Block Feedback | Trial Feedback |
| --- | --- | --- | --- | --- | --- | --- | --- |
| Barker (2015) | Volunteer | LSAS-SR | Flanker | Arrow | Two-handed | With | Without |
| Beste (2013) | Volunteer | ASI | Flanker | Arrow | Two-handed | Without | With |
| Boksem (2006) | Volunteer | BIS | Flanker | Letter | Two-handed | Without | Without |
| Carrasco (2013a) | Clinical - pediatric OCD | K-SADS-PL | Flanker | Arrow | Two-handed | With | Without |
| Carrasco (2013b) | Clinical - pediatric OCD | K-SADS-PL | Flanker | Arrow | Two-handed | With | Without |
| Cavanagh (2008) | Volunteer | BIS | Flanker | Letter | Two-handed | Without | With |
| Chang (2010) | Volunteer | ASR | Flanker | Letter | Two-handed | Without | Without |
| Endrass (2014) | Clinical-OCD/SAD | SCID | Flanker | Arrow | Two-handed | With | Without |
| Grützmann (2016) | Clinical - OCD | SCID | Flanker | Arrow | Two-handed | With | Without |
| Hanna (2012) | Clinical -pediatric OCD | K-SADS-PL | Flanker | Arrow | Two-handed | With | Without |
| Hanna (2016) | Clinical - pediatric OCD | Existing diagnosis | Flanker | Arrow | Two-handed | With | Without |
| Hanna (2018) | Clinical – adolescent OCD | K-SADS-PL | Flanker | Arrow | Two-handed | With | Without |
| Kaczurkin (2013) | Volunteer | OCI-R | Flanker | Letter |  | Without | With |
| Klawohn (2014) | Clinical - OCD | SCID | Flanker | Arrow | Two-handed | With | Without |
| Klawohn (2016) | Clinical - OCD | SCID | Modified Flanker | Arrow | Two-handed | With | Without |
| Ladouceur (2018) | Clinical – pediatric anxiety | K-SADS-PL | Flanker | Arrow | Two-handed | With | Without |
| Larson & Clayton (2011)* | Volunteer | STAI-T | Flanker | Arrow | One-handed | Without | Without |
| Larson (2011)* | Volunteer | STAI-T | Flanker | Arrow | One-handed | Without | Without |
| Larson (2013)* | Clinical – GAD | SCID | Flanker | Letter | One-handed | Without | Without |
| Liu (2014) | Clinical – pediatric OCD | K-SADS-PL | Flanker | Arrow | Two-handed | With | Without |
| Luu (2000) | Volunteer | PANAS | Flanker | Letter | Two-handed | Without | With |
| Meyer (2012) | Volunteer - child | Parent – SCARED | Flanker | Arrow | One-handed | With | Without |
| Meyer (2017) | Volunteer | STAI-T | Flanker | Arrow | One-handed | With | Without |
| Milyavskaya (2018)* | Volunteer | BIS | Flanker | Arrow | Two-handed | Without | Without |
| Moran (2012) | Volunteer | PSWQ | Flanker | Letter | One-handed | Without | Without |
| Olvet (2009) | Volunteer | DASS | Flanker | Arrow | One-handed | With | Without |
| Olvet (2012) | Clinical - PTSD | BFI-N | Flanker | Arrow | One-handed | With | Without |
| Rabinak (2013) | Clinical - OCD | SCID | Flanker | Arrow | One-handed | With | Without |
| Riesel (2011) | Clinical - OCD | SCID | Flanker | Arrow | Two-handed | With | Without |
| Riesel (2014) | Clinical - OCD | SCID | Flanker | Arrow | Two-handed | With | Without |
| Riesel (2015) | Clinical - OCD | SCID | Flanker | Arrow | Two-handed | With | Without |
| Roh (2016) | Clinical - OCD | SCID | Modified Flanker | Arrow | Two-handed | With | Without |
| Roh (2017) | Volunteer – child | SCID | Modified Flanker | Arrow | Two-handed | With | Without |
| Santesso (2005) | Volunteer - child | JEPQR-S | Flanker | Letter | Two-handed | Without | Without |
| Santesso (2006) | Volunteer - child | CBCL | Flanker | Letter | Two-handed | Without | Without |
| Santesso (2009) | Volunteer – child | Composite | Flanker | Letter | Two-handed | Without | Without |
| South (2010) | Clinical - OCD | BIS | Flanker | Arrow | One-handed | Without | Without |
| Stern (2010) | Clinical - PTSD | SCID | Modified Flanker | Letter | Two-handed | With | With |
| Swick (2015) study 1 | Clinical - PTSD | PCL-M | Flanker | Arrow | Two-handed | Without | Without |
| Swick (2015) study 2 | Volunteer | PCL-M | Modified Flanker | Arrow | Two-handed | Without | Without |
| Tanovic (2017) | Volunteer | PSWQ | Flanker | Arrow | Two-handed | With | Without |
| Tops (2011) | Clinical - GAD | BIS | Flanker | Letter | Two-handed | Without | Without |
| Weinberg (2010) | Clinical - GAD | SCID | Flanker | Arrow | One-handed | With | Without |
| Weinberg (2012) | Clinical | SCID | Flanker | Arrow | One-handed | With | Without |
| Weinberg (2015) | Clinical – GAD/OCD | SCID | Flanker | Arrow | One-handed | With | Without |
| Xiao (2011) | Clinical – GAD/OCD | Chinese MINI | Flanker | Letter | Two-handed | Without | With |

Note: \* denotes unpublished study. GAD, generalized anxiety disorder; OCD, obsessive-compulsive disorder; PTSD, post-traumatic stress disorder. ASR, Achenbach Self-Report; BFI-N, Big Five Inventory-Neuroticism; BIS, Behavioral Inhibition System Scale; CBCL, Child Behavior Checklist; DASS, Depression Anxiety Stress Scale; JEPQR-S, Junior Eysenck Personality Questionnaire – Revised, Self-report; K-SADS-PL, Schedule for Affective Disorders and Schizophrenia for School-Age Children-Present and Lifetime Version; LSAS-SR, Liebowitz Social Anxiety Scale – Self-Report; MINI, Mini-International Neuropsychiatric Interview; PANAS, Positive and Negative Affect Schedule; PCL-M, PTSD Checklist, Military version; PSWQ, Penn State Worry Questionnaire; SCARED, Screen for Child Anxiety Related Disorders; SCID, Structured Clinical Interview for DSM Disorders; STAI-T, State and Trait Anxiety Inventory-Trait Version.

Consistent with prior work by Moser and colleagues (2013), Saunders and Inzlicht (2020) also coded whether studies employed a measure of anxiety that mapped most closely onto the worry/anxious apprehension component of anxiety, or if the anxiety measure reflected a broader mix of anxiety dimensions. To maximize the number of included studies, for all primary analyses reported in the main text we included studies regardless of the anxiety measure used. However, we also performed an exploratory set of meta-regression analyses testing for moderation restricted to only studies employing anxiety measures that mapped onto the anxious apprehension/worry component of anxiety and obtained similar results (see supplement).

### Data extraction

#### Effect sizes

Given that effect sizes employed in the meta-analysis were drawn directly from the Saunders and Inzlicht (2020) database, we provide a brief description of the procedures used by Saunders and Inzlicht (2020); for complete details, readers should refer to the original paper. For all studies, Saunders and Inzlicht (2020) extracted effect sizes associated with the ERN on error trials. In cases where studies employed a manipulation of motivation/valence, the effect size was extracted from the neutral condition. Note that for a minority of studies, effect sizes associated with the delta-ERN were also extracted; however, there were not enough to allow for moderation analyses. Thus, our moderation analyses all focus on associations between anxiety and the ERN on error trials. For studies employing continuous measures of anxiety, Saunders and Inzlicht (2020) extracted Pearson’s *r*; when one or more measures of anxiety were reported, the effect size was extracted for the analysis most closely mapping onto the construct of worry/anxious apprehension, consistent with prior work by Moser and colleagues (2013). For studies involving group comparison, Saunders and Inzlicht (2020) computed Cohen’s *d* and then converted the result into Pearson’s *r*. Effect sizes were only included if drawn from independent samples (Saunders & Inzlicht, 2020). For complete details on effect size extraction, see: Saunders and Inzlicht (2020).

#### Extraction and coding of moderator variables

We coded the following variables for all Flanker task studies selected for inclusion: 1) stimulus type (arrow, letter), 2) response type (one-handed, two-handed), and 3) block-level feedback (with, without). Note that for response type, a one-handed response format implies responding using separate fingers on the same hand. To isolate any potential moderating role of block-level feedback, we additionally coded whether trial-level feedback was also presented and removed any such studies (n = 6) from analyses testing the moderating role of block-level feedback. Note that we did not attempt to directly assess the effect of trial-level feedback on its own as a moderator, given the low number of studies employing trial-level feedback.

For published studies, two independent coders extracted/coded the variables of interest. Extracted values were then reconciled to ensure complete agreement between both coders. In cases where agreement could not be reached or the original paper was missing necessary information, authors were contacted via email. Note that we coded studies as being “without” block-level feedback in cases where the original article did not mention block-level feedback, and then attempted to confirm these codes with the authors; if the authors provided alternative information, the codes were updated, otherwise the “without” code was retained (for 6 studies, we were unable to confirm with the authors). For unpublished studies, relevant information was obtained from the Saunders and Inzlicht (2020) database and authors of the original studies were emailed to request missing task parameter information. First and senior authors were contacted up to two times, with a final response rate of 87.88%. Following these procedures, out of the 46 flanker tasks selected for inclusion, all 46 had viable stimulus type information, 45 had viable response type information, and 40 had viable block-level feedback information (after removing 6 studies that also included trial-level feedback).

### Meta-Analyses

To serve as a baseline for subsequent tests of moderation, we first fit a random-effects meta-analysis model to estimate the overall effect size of ERN-anxiety associations amongst Flanker task studies, as well as to estimate total heterogeneity via I^2^. Next, we fit a series of three random-effects meta-regression models to test whether stimulus type, response type, or block-level feedback moderated the effect size associated with ERN-anxiety associations amongst Flanker task studies. All analyses employed the *metafor* package in R (Viechtbauer & Viechtbauer, 2015). Effect sizes (Pearson’s *r*) were first transformed into Fisher’s Z via the *escalc* function (Viechtbauer & Viechtbauer, 2015). For each moderation model, moderator significance was assessed via the Q test and estimates of effect size at each level of the moderator were predicted based on model coefficients derived from the same model. Restricted maximum-likelihood estimation (REML) was used in all meta-analytic estimates of effect size.

For each moderation model, we assessed publication bias visually via the construction of a funnel plot (Sterne & Egger, 2001), as well as quantitatively via Egger’s test (Egger et al., 1997). In cases where Egger’s test was significant for a given moderation model, we further assessed publication bias (via Egger’s test) in separate models fit at each level of the moderator (subgroup analysis) and estimated the bias-corrected effect size for each subgroup via the trim-and-fill procedure (Duval & Tweedie, 2000).

## Results

### Overall effect size for Flanker task studies

Amongst all Flanker task studies investigated, the overall effect size for the ERN-anxiety association was significant and estimated as -0.208 (95% CI = -0.261, -0.154), meaning a more negative (larger) ERN amplitude was associated with greater anxiety symptom severity. Total heterogeneity was significant, Q(45) = 93.18, *p* < .001, I^2^ = 51.55%, suggesting ample variability across studies to support further moderation analyses. These results served as a baseline for subsequent analyses of moderation.

### Stimulus type

Stimulus type did not significantly moderate the effect size of ERN-anxiety associations, Q(1) = 0.011, *p* = .916. The predicted effect sizes were similar for studies with arrow stimuli, -.206 (95% CI: -0.270, -0.143), and letter stimuli, -0.213 (95% CI: -0.318, -0.108); see Figure 1 and Table 2. Egger’s test of publication bias was not significant (*p* = .326), suggesting there was no evidence of publication bias amongst studies employed to test the moderating role of stimulus type; see Figure 2.

**Figure 1.**
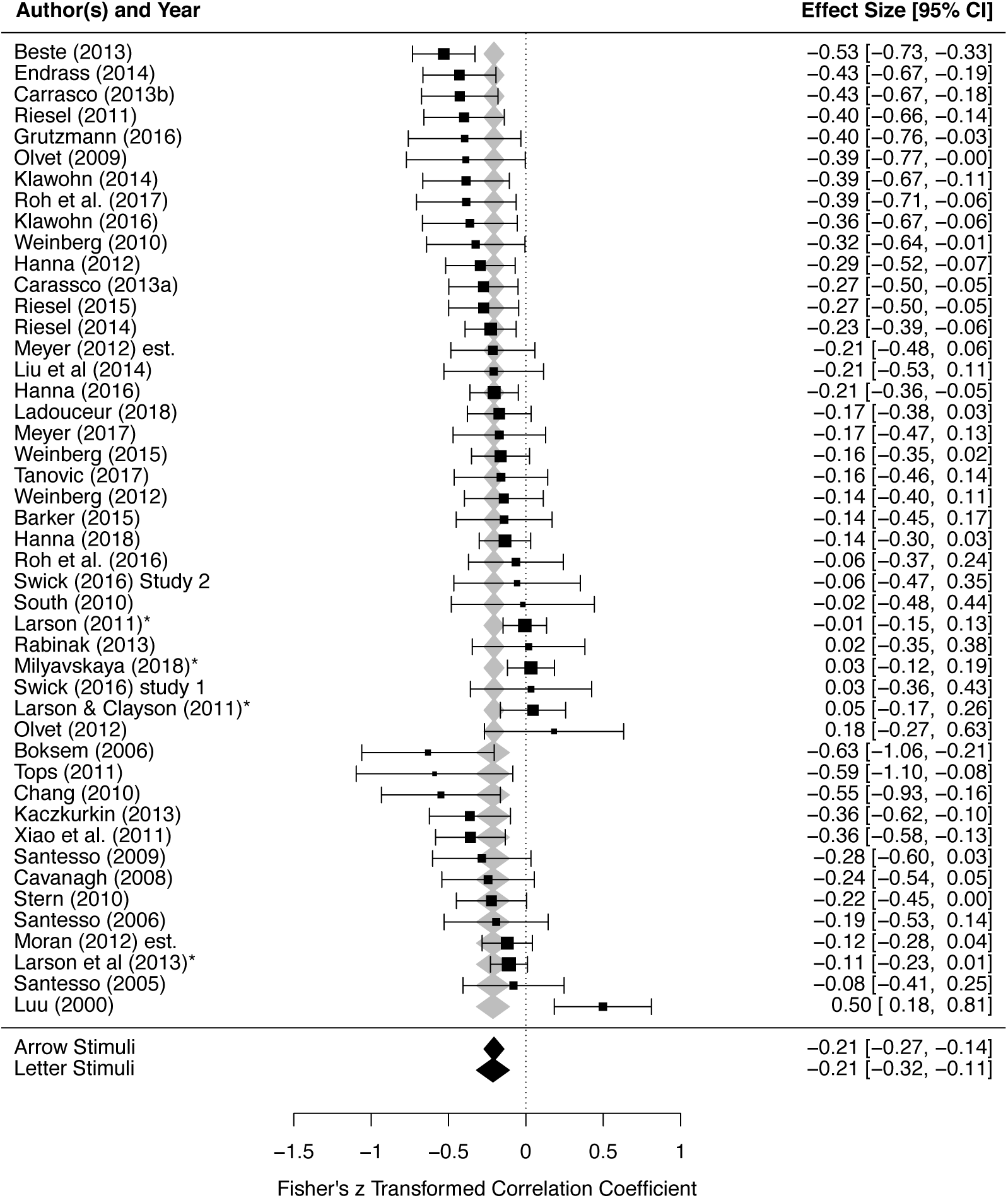
Stimulus type does not significantly moderate the effect sizes of ERN-anxiety associations. Order of studies are grouped by stimulus type (Arrow followed by Letter), and further ordered by each study’s effect size within each group. Each study’s effect size, as well as weight within the meta-analysis, is depicted via the horizontal position and size of black boxes, respectively; 95% CI for each study’s effect size is depicted via the length of horizontal lines extending from each black box. Predicted average effect sizes and their 95% CI, at each level of the moderator, are depicted in terms of the horizontal position and width of the diamonds, respectively (gray/black diamonds are redundant). ERN, error-related negativity.

**Figure 2.**
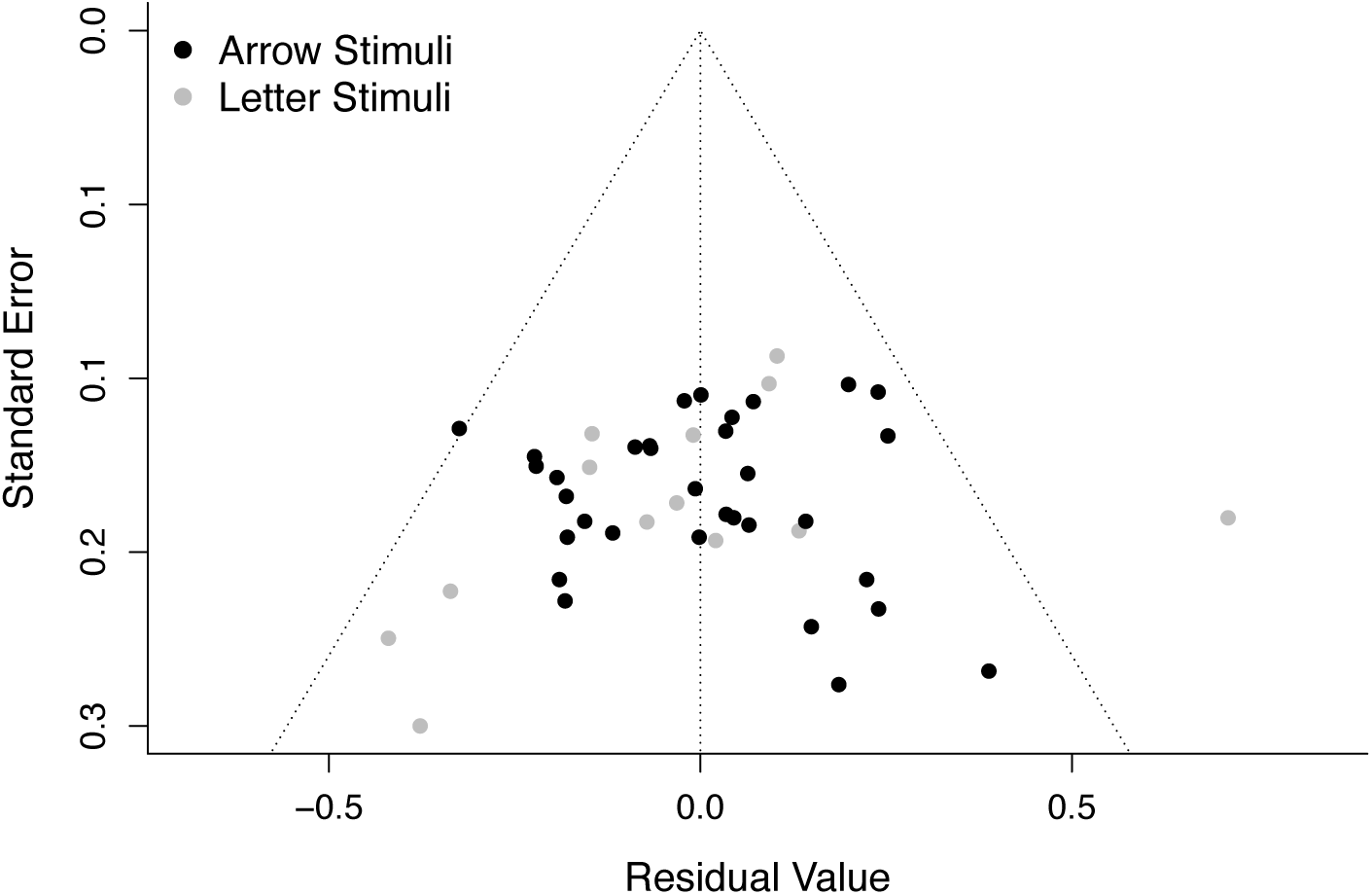
Funnel plot depicting no evidence of publication bias amongst the studies employed to test the moderating role of stimulus type on ERN-anxiety associations. Standard errors are plotted as a function of residual values, with each individual study depicted as a single point. Black and gray points denote studies that employed arrow vs. letter stimuli, respectively. Note that Egger’s test of publication bias was not significant (*p* = .326), suggesting there was no evidence of publication bias. ERN, error-related negativity.

**Table 2.** Effect of task parameters on effect sizes associated with ERN-anxiety associations.

|  | **Q** | **p** | **effect** | **95% CI** | **K** | **N** |
| --- | --- | --- | --- | --- | --- | --- |
| **Stimulus type** | 0.011 (1) | 0.916 |  |  |  |  |
| **Arrow** |  |  | -0.206 | -0.270, -0.143 | 33 | 2342 |
| **Letter** |  |  | -0.213 | -0.318, -0.108 | 13 | 913 |
| **Response type** | 5.861 (1) | 0.016* |  |  |  |  |
| **One-handed** |  |  | -0.108 | -0.200, -0.016 | 13 | 1128 |
| **Two-handed** |  |  | -0.244 | -0.305, -0.184 | 32 | 2068 |
| **Block-level feedback** | 14.585 (1) | <.001* |  |  |  |  |
| **Without** |  |  | -0.088 | -0.147, -0.029 | 14 | 1140 |
| **With** |  |  | -0.237 | -0.285, -0.189 | 26 | 1714 |

Note: \*denotes significance. ERN, error-related negativity.

### Response type

Response type significantly moderated the effect size of ERN-anxiety associations, Q(1) = 5.861, *p* = .016. The predicted effect size was larger (more negative) for two-handed studies, - 0.244 (95% CI: -0.305, -0.184), compared to one-handed studies, -0.108 (95% CI: -0.200, - 0.016); see Figure 3 and Table 2. Egger’s test of publication bias was not significant (*p* = .441), suggesting there was no evidence of publication bias amongst studies employed to test the moderating role of response type; see Figure 4.

**Figure 3.**
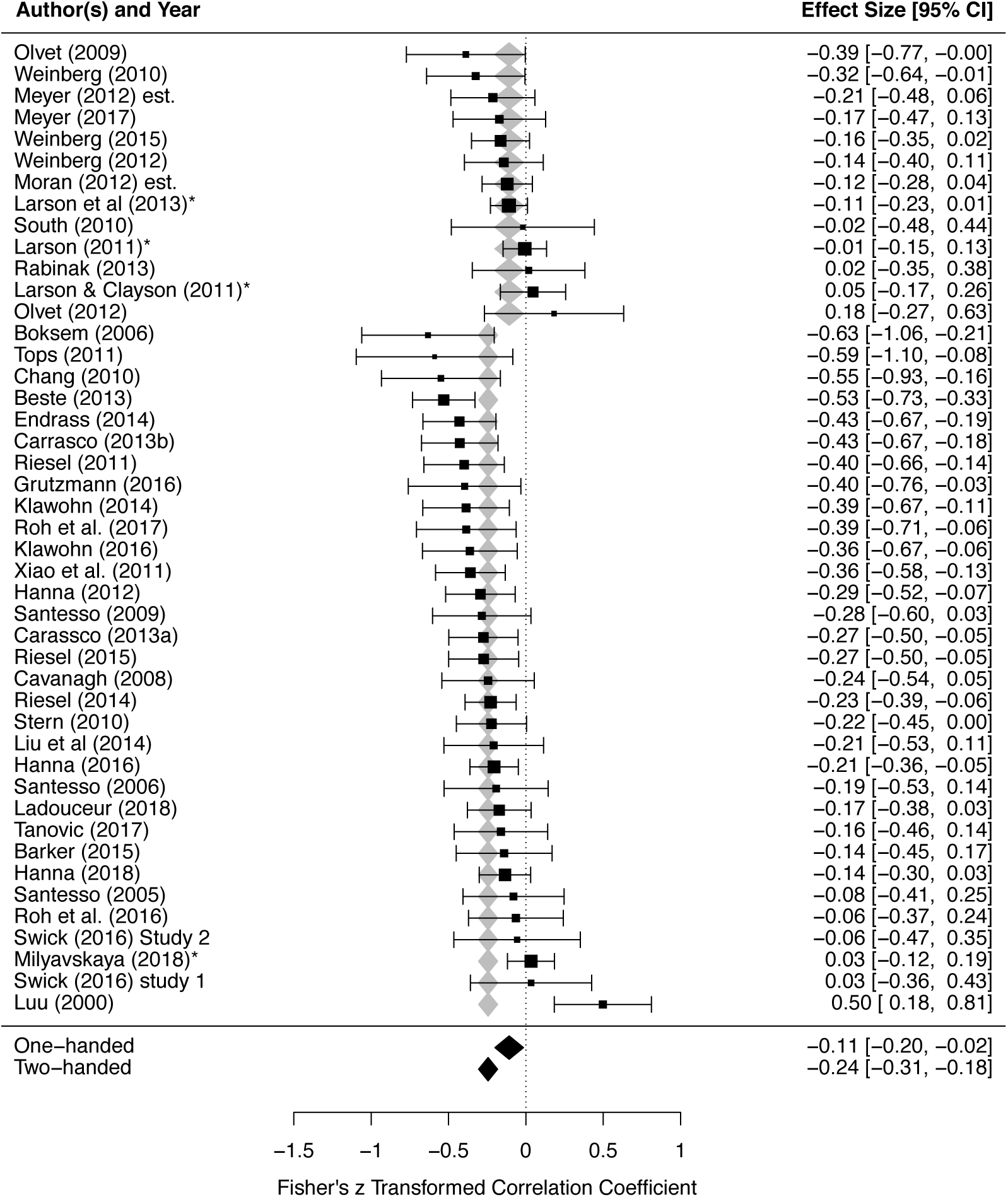
Response type significantly moderates the effect size of ERN-anxiety associations. Order of studies are grouped by response type (one-handed followed by two-handed), and further ordered by each study’s effect size within each group. Each study’s effect size, as well as weight within the meta-analysis, is depicted via the horizontal position and size of black boxes, respectively; 95% CI for each study’s effect size is depicted via the length of horizontal lines extending from each black box. Predicted average effect sizes and their 95% CI, at each level of the moderator, are depicted in terms of the horizontal position and width of the diamonds, respectively (gray/black diamonds are redundant). ERN, error-related negativity.

**Figure 4.**
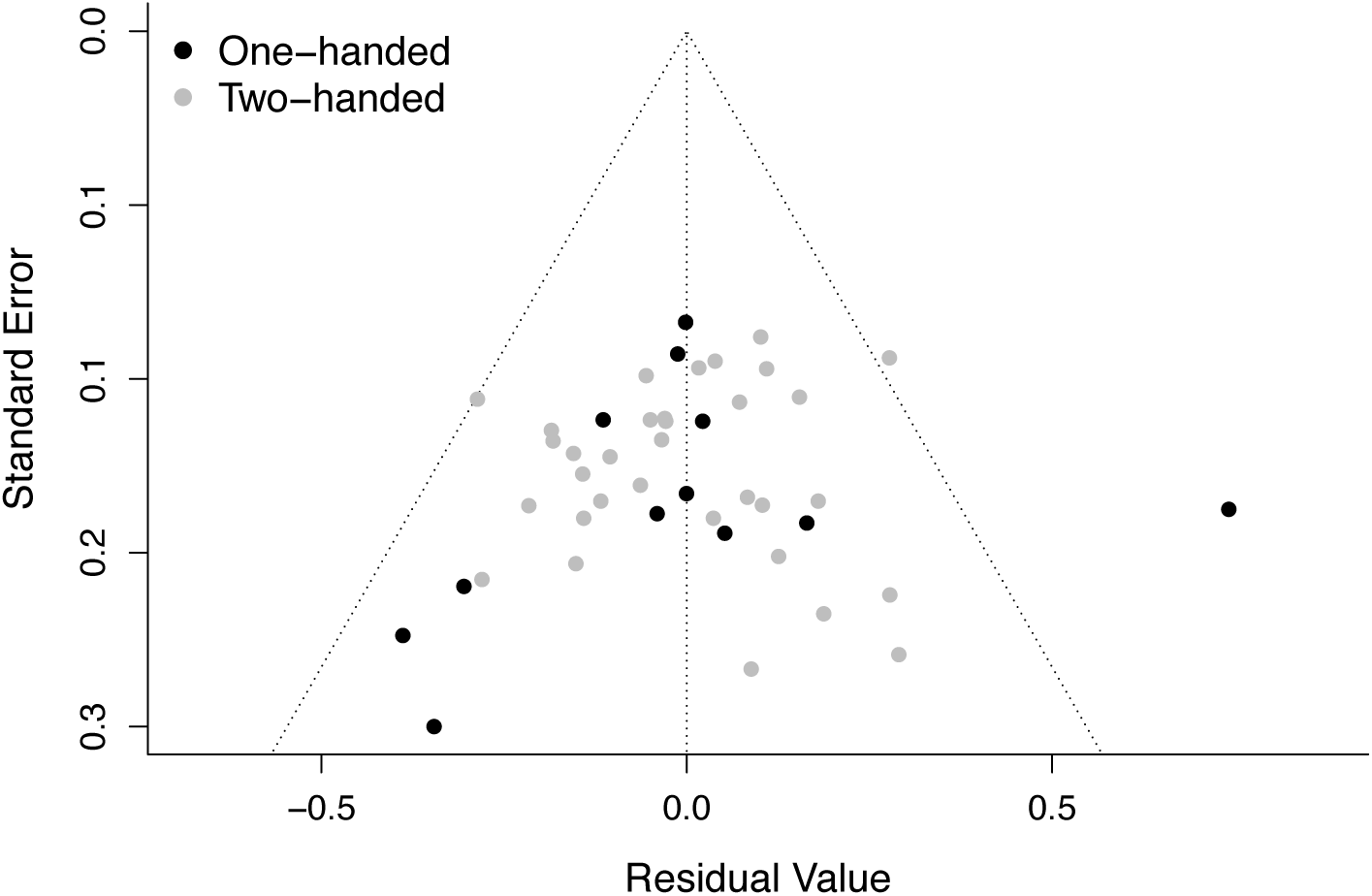
Funnel plot depicting no evidence of publication bias amongst the studies employed to test the moderating role of response type on ERN-anxiety associations. Standard errors are plotted as a function of residual values, with each individual study depicted as a single point. Black and gray points denote one-handed vs. two-handed studies, respectively. Note that Egger’s test of publication bias was not significant (*p* = .441), suggesting there was no evidence of publication bias. ERN, error-related negativity.

### Block-level feedback

Block-level feedback significantly moderated ERN-anxiety associations, Q(1) = 14.585, *p* < .001. The predicted effect size was larger (more negative) for studies with block-level feedback, -0.237 (95% CI: -0.285, -0.189), compared to studies without block-level feedback, - 0.088 (95% CI: -0.147, -0.029); see Figure 5 and Table 2.

**Figure 5.**
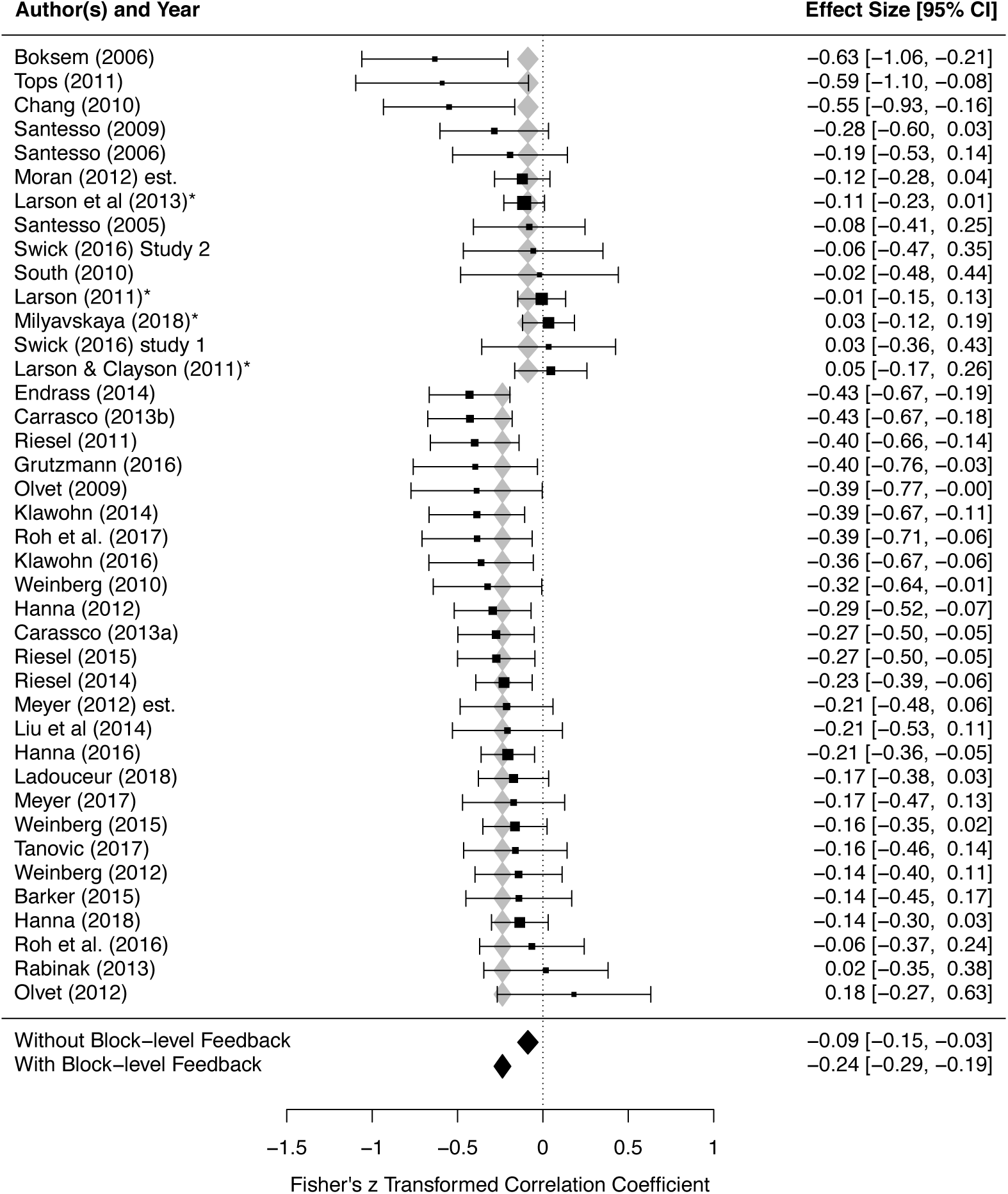
Block-level feedback significantly moderates the effect size of ERN-anxiety Order of studies are grouped by whether studies included block-level feedback (without followed by with), and further ordered by each study’s effect size within each group. Each study’s effect size, as well as weight within the meta-analysis, is depicted via the horizontal position and size of black boxes, respectively; 95% CI for each study’s effect size is depicted via the length of horizontal lines extending from each black box. Predicted average effect sizes and their 95% CI, at each level of the moderator, are depicted in terms of the horizontal position and width of the diamonds, respectively (gray/black diamonds are redundant). ERN, error-related negativity.

Egger’s test of publication bias was significant (*p* = 0.030), consistent with possible publication bias amongst studies employed to test the moderating role of block-level feedback; see Figure 6. Thus, we further assessed publication bias (via Egger’s test) in separate models fit at each level of the moderator (subgroup analysis) and estimated the bias-corrected effect size for each subgroup via the trim-and-fill procedure (see supplement) (Duval & Tweedie, 2000). Results of these additional analyses demonstrated evidence of publication bias only among studies without block-level feedback. Correcting for bias via the trim-and-fill procedure yielded a non-significant effect size estimate among studies without block-level feedback, whereas the effect size for studies with block-level feedback remained significant (see supplement).

**Figure 6.**
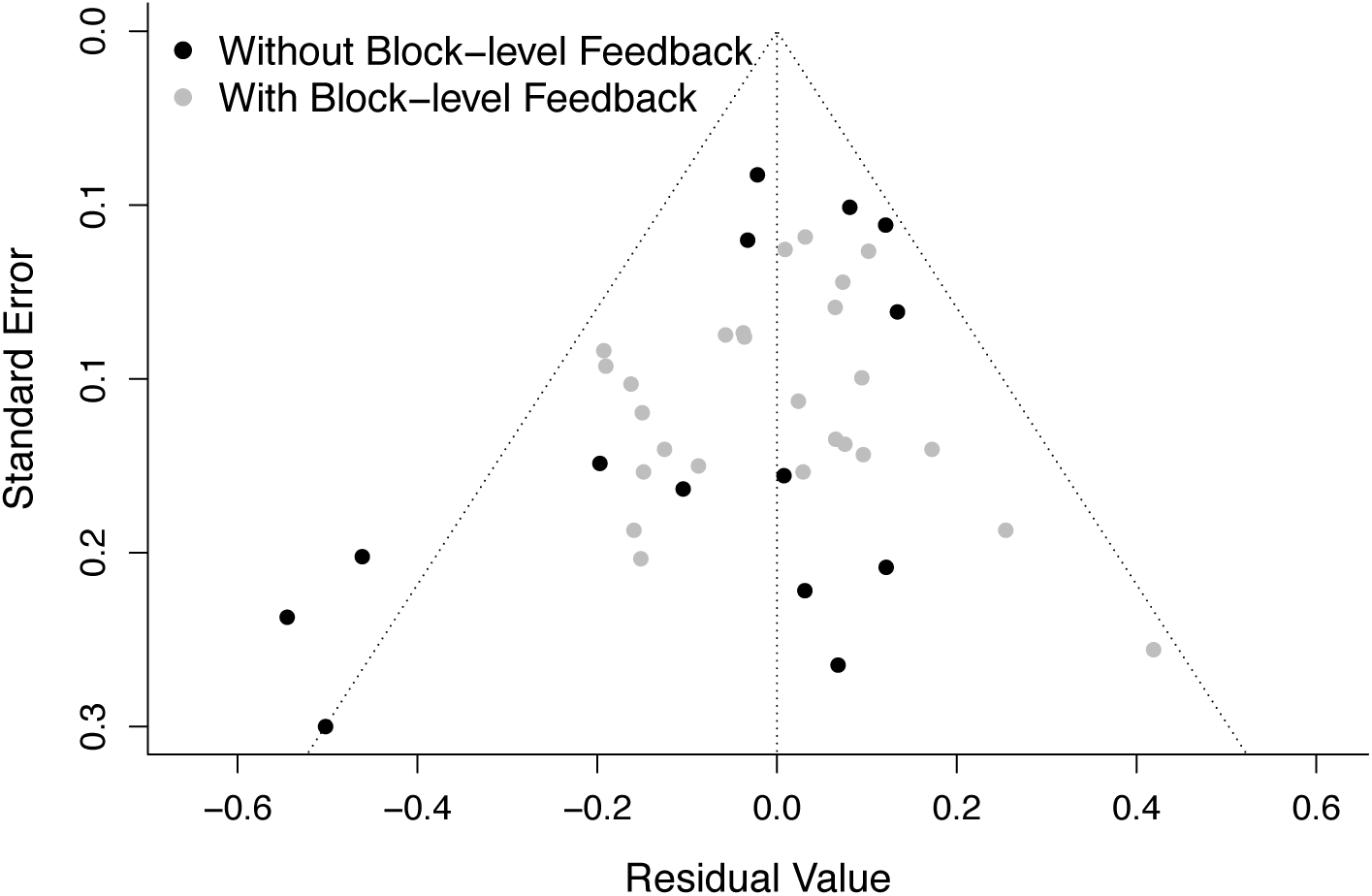
Funnel plot suggesting possible publication bias amongst the studies employed to test the moderating role of block-level feedback on ERN-anxiety associations. Standard errors are plotted as a function of residual values, with each individual study depicted as a single dot. Black and gray points denote studies without vs. with block-level feedback, respectively. Note that Egger’s test of publication bias was significant (*p* = 0.030), consistent with possible publication bias. Further subgroup analyses suggested that possible publication bias was exclusive to studies without block-level feedback, not those with block-level feedback (see main text and supplement for further details). ERN, error-related negativity.

## Discussion

The goal of this study was to determine whether variation in several Flanker task parameters (stimulus type, response type, and block-level feedback) is associated with differences in the magnitude of ERN-anxiety associations. To this end, we did not find evidence that stimulus type (arrow vs. letter) impacted ERN-anxiety associations. However, we found that both response type and block-level feedback significantly moderated ERN-anxiety associations. ERN-anxiety associations were strongest for two-handed (vs. one-handed) tasks, and for tasks with (vs. without) block level feedback. Given that these task parameters commonly vary across studies investigating ERN-anxiety associations, the presence of such moderating effects could, at least in part, explain some inconsistencies in reported effect sizes in the published literature and across prior meta-analyses. Moreover, the results of this meta-analysis can directly inform the design of studies seeking to maximize ERN-anxiety associations. Below, we discuss practical implications of these findings, followed by a more in-depth discussion of their potential theoretical implications.

### Practical implications

At a practical level, the current results emphasize that variation in Flanker task parameters can influence the magnitude of ERN-anxiety associations for a given study. Of note, both response type and block-level feedback were not only found to have a significant impact on ERN-anxiety associations, but the reported details of these parameters were absent from at least some published studies (requiring us to reach out to authors to obtain this information). Given that our results suggest that these task parameters may influence the magnitude of ERN-anxiety associations, we encourage researchers to include such details in future publications.

The results can also serve to provide guidance on the set of task parameters that can be used to maximize ERN-anxiety associations: a two-handed Flanker task with block-level feedback. It is worth noting that such a design affords further benefits. Two-handed tasks allow for performing further analyses as a function of response hand, crucial for testing at least one prominent theory of ERN-anxiety associations (CEMH; Lin et al., 2015; Moser et al., 2013).

Moreover, two-handed tasks allow for computing measures of effector-specific motor activity (e.g., the lateralized readiness potential or lateralized beta; Fischer et al., 2018; Smulders et al., 2012), providing further insight into the dynamics of error monitoring. Presenting block-level feedback that is contingent on behavioral performance (consistent with the recommendations of Gehring et al., 2012) has the added benefit of ensuring adequate error rates for reliable ERN analyses. As we discuss further below, it is worth noting that including performance-contingent feedback also implies that the type of feedback presented is not uniform across participants, which could introduce further between-subject variance (Gloe & Louis, 2021). Nonetheless, until further data demonstrates this to be problematic, we recommend including block-level feedback for studies seeking to maximize ERN-anxiety associations. Future studies are needed to further examine exactly why block-level feedback and two-handed tasks increase ERN-anxiety associations. Additionally, work is needed to understand how other parameters may impact ERN-anxiety associations and/or interact with response type and block-level feedback. However, based on extant data, and if the primary goal is to maximize ERN-anxiety associations, then employing a two-handed Flanker task with block-level feedback is recommended.

### Theoretical implications

#### Differences by response type

We observed a significant effect of response type on the magnitude of ERN-anxiety associations; however, the theoretical explanation for this phenomenon remains undetermined, as multiple theories can account for the effect. One possible explanation is provided by comparator/mismatch theories of the ERN (Bernstein et al., 1995; Coles et al., 2001; Falkenstein et al., 1996). As noted in the introduction, there is a longstanding debate over whether the ERN arises from conflict monitoring (Botvinick et al., 2001; Yeung et al., 2004) vs. a comparator/mismatch detection process (Bernstein et al., 1995; Coles et al., 2001; Falkenstein et al., 1996), among other possible explanations of the ERN. Conflict monitoring theory proposes that ERN magnitude reflects the degree of conflict—formalized as Hopfield energy (Hopfield, 1982)—between simultaneously active correct and incorrect responses (Botvinick et al., 2001; Yeung et al., 2004). One interpretation of conflict monitoring theory (Gehring & Fencsik, 2001) is that more similar motor responses should elicit greater conflict and a larger ERN. In contrast, comparator/mismatch theories propose the ERN arises from a mismatch detection process in which the actual (error) motor response is compared with the correct motor response (derived from continued stimulus processing; Coles et al., 2001). Crucially, comparator/mismatch theories predict a larger ERN for tasks involving more differentiated motor responses (greater possible mismatch between error/correct motor responses (Bernstein et al., 1995; Coles et al., 2001; Falkenstein et al., 1996). Ultimately, the ERN may arise from both conflict monitoring and comparator/detection processes (Arbel & Donchin, 2011; Bernstein et al., 1995; Falkenstein et al., 1996; Gehring & Fencsik, 2001).

Nonetheless, the degree to which the ERN elicited by a given task is driven by conflict vs. a comparator/detection process should differ as a function of how similar the motor responses are for that task. Moreover, to the extent that anxiety is more closely associated with ERN variation arising from conflict vs. a comparator/detection process, differences in the magnitude of ERN-anxiety associations should also emerge across tasks as a function of how similar the motor responses are within each task. Consistent with the notion that motor commands are organized hierarchically, prior work demonstrates that preparation of motor responses for two different fingers on the same hand involve more similar motor commands as opposed to fingers on opposite hands (Miller, 1982; see also: Rosenbaum, 1980). Thus, our finding that ERN-anxiety associations were stronger for two-handed tasks (presumably involving more differentiated motor responses and an ERN more strongly driven by comparator/mismatch detection) provides indirect evidence against the notion that ERN-anxiety associations are driven by ERN variation associated with conflict monitoring. Instead, our results are more consistent with the notion that ERN variation arising from a comparator/mismatch process is what more closely relates to anxiety. Nonetheless, it is important to note that this conclusion draws on the assumption that similarity of motor responses can be used to differentiate between conflict monitoring and comparator/detection theories of the ERN (Gehring & Fencsik, 2001). Thus, a more parsimonious interpretation of these results is simply that ERN-anxiety associations are stronger for tasks with more differentiated motor responses. More importantly, alternative theories can also explain the effect of response type on ERN-anxiety associations, including the CEMH theory (Lin et al., 2015; Moser et al., 2013) discussed in the following section. Thus, to provide a more direct test of whether ERN-anxiety associations are driven by ERN variation associated with a comparator/mismatch detection process, experimental studies are needed that assess ERN-anxiety associations when the similarity of motor response options are parametrically modulated at a within-subjects level. This would allow for assessing whether parametric decreases in response similarity predict commensurate increases in ERN-anxiety associations.

The finding that ERN-anxiety associations were stronger for two-handed tasks is also consistent with the CEMH theory (Lin et al., 2015; Moser et al., 2013), which states that ERN-anxiety associations are driven by anxious apprehension/worry creating asymmetries in the relative prepotency of left-vs. right-hand responses. To understand this prediction, it is necessary to first review the logic of the CEMH theory. The CEMH theory (Lin et al., 2015; Moser et al., 2013) draws on work by Hochman (Hochman et al., 2014), which interprets the ERN as reflecting not a detection process, but a regulative process reflecting inhibition of an incorrect response to allow for a corrective response. Within this view, greater inhibition is required (and a larger ERN observed) when an error response is relatively more prepotent than the correct response (stronger activation needed to inhibit the incorrect response) and when the error response shares fewer motor commands with the correct response (more incorrect motor command features to inhibit; Hochman et al., 2014). Drawing on these ideas, Moser and colleagues (Lin et al., 2015; Moser et al., 2013) propose that, within a two-handed response task, greater verbal rumination—associated with anxious apprehension/worry—primes left motor cortex (Engels et al., 2007; Oathes et al., 2008), in turn increasing the relative prepotency of right-vs. left-hand responses (Lin et al., 2015). As a result, when high anxious individuals perform a two-handed task, errors made with the right hand are more prepotent and require greater inhibition (eliciting a larger ERN; Hochman et al., 2014; Lin et al., 2015). Thus, ERN-anxiety associations are proposed to be driven by right-handed responses made within the context of a two-handed task. Confirming these predictions, in two studies employing two-handed flanker tasks, ERN-anxiety associations were only present for right-handed responses (Lin et al., 2015). Here, it is important to emphasize that in addition to proposing ERN-anxiety associations are driven by right-hand responses, the CEMH theory also implicitly assumes that this is only possible within a two-handed task. In other words, the CEMH would not predict stronger ERN-anxiety associations in a one-handed task, even if all responses were made by the right hand. This is because only in a two-handed task can left-lateralized verbal rumination lead to a *relative* increase (difference) in the prepotency between right-(vs. left-) hand responses. Therefore, our finding that ERN-anxiety associations were stronger in the two-handed (vs. one-handed) task is fully consistent with the CEMH theory.

Although the finding that ERN-anxiety associations were stronger in two-handed tasks is consistent with the predictions of the CEMH theory (Lin et al., 2015; Moser et al., 2013), other theories can also account for this effect, and one of our supplementary analyses was at least partially inconsistent with the CEMH theory. Our primary analyses reported in the main text all focus on associations between the ERN and anxiety, broadly defined (i.e., without distinguishing between worry/anxious apprehension and other dimensions of anxiety). However, within the supplement, we report the results of a follow-up analysis that assessed the moderating role of response type when only including studies employing anxiety measures mapping onto anxious apprehension/worry (see supplement). The CEMH theory would predict that the effect of response type on the magnitude of ERN-anxiety associations would be at least as strong, if not stronger, when restricted to only studies employing measures of anxious apprehension/worry. However, this was not the case, with the effect of response type being only marginal (*p* = .061) in the analysis restricted to studies assessing anxious apprehension/worry (although we urge caution interpreting these results, as this difference could be attributed to differences in statistical power). Regardless, future studies are needed to directly test whether the effect of response type is best explained by the CEMH or alternative theories. For example, a more direct test would come from a study in which each participant performs both a one and a two-handed version of the Flanker. This would allow for not only replicating the effect of response type on ERN-anxiety associations at the within-subject level, but also allow for testing whether any increase in ERN-anxiety associations for the two-handed task is fully mediated by right-handed responses.

#### Differences by block-level feedback

Evidence of possible publication bias for studies included in the analysis of block-level feedback does not appear to explain the finding that ERN-anxiety associations were stronger for studies with block-level feedback (vs. without). Subsequent subgroup analyses (see supplement) found no evidence of bias among studies with block-level feedback, only among studies without block-level feedback. Moreover, correcting for bias only served to produce a non-significant effect size estimate among studies without block-level feedback, whereas the effect size for studies with block-level feedback remained significant following correction (see supplement). Analyses of publication bias are imperfect and not without limitations (Thornton & Lee, 2000). Nonetheless, the results raise the possibility that not only are ERN-anxiety associations stronger amongst studies with block-level feedback, but that studies without block-level feedback may be underreported in the published literature.

At a theoretical level, multiple theories can explain the finding that ERN-anxiety associations were stronger amongst studies with block-level feedback. One possibility is that presenting block-level feedback of any form serves to remind participants that their performance is being monitored/evaluated by researchers, serving as a quasi-social-evaluative context manipulation, similar to explicit manipulations of social context (e.g., Barker et al., 2015, 2018; Buzzell et al., 2019; Hajcak et al., 2005). Importantly, ERN associations with social anxiety and related symptoms are stronger within a social-evaluative context (Barker et al., 2015; Buzzell et al., 2017; Niu et al., 2023). Our meta-analysis conceptualized anxiety broadly, consistent with prior work (Cavanagh & Shackman, 2015; Moser et al., 2013; Saunders & Inzlicht, 2020). To the extent that broad measures of anxiety also capture social anxiety and related symptoms, one hypothesis is that stronger ERN-anxiety associations in studies with block-level feedback result from increased social-evaluative concerns, particularly for individuals high in social anxiety or related symptoms. Consistent with theories that posit the ERN-anxiety association is driven by generalized defensive reactivity to threat (for a review, see: Weinberg et al., 2012), a related possibility is that presenting block-level feedback does not create a social-evaluative context per se, but nonetheless creates a situation of increased “pressure” that induces a stronger defensive response and concomitant increase in the ERN. To test these related—but distinct—hypotheses, a study is needed that directly manipulates the presence of block-level feedback and assesses broadband anxiety as well as social anxiety and related symptoms. Both theories would predict that the presence of block-level feedback would increase ERN-anxiety associations. However, theories of defensive reactivity (Weinberg, Riesel, et al., 2012) would predict that the effect of block-level feedback on ERN-anxiety associations would be best explained by variation in broadband and/or general anxiety measures. In contrast, the social-evaluative hypothesis would predict that the effect of block-level feedback on ERN-anxiety associations would be best explained by variation in social anxiety or related symptoms; based on recent work, the construct of Fear of Negative Evaluation may be especially prominent in explaining such ERN-anxiety associations (Niu et al., 2023).

An additional possibility is that the effect of block-level feedback on ERN-anxiety associations arises from differences in how anxious vs. non-anxious individuals modify their error monitoring in response to feedback. To this end, a prior study by Riesel and colleagues (2019) manipulated whether participants performed a task within the context of instructions emphasizing either speed or accuracy, along with associated trial-*and* block-level feedback. Under speed (vs. accuracy) instructions, non-anxious individuals showed a substantial reduction in their ERN; conversely, individuals diagnosed with OCD showed a much more modest reduction in their ERN (Riesel et al., 2019). Thus, it is possible that anxious and non-anxious differ in the degree to which they adjust error monitoring levels in response to speed-related feedback. Non-anxious individuals may be more able to modulate levels of error monitoring in accord with task instructions, resulting in a reduced ERN within the context of speed (vs. accuracy) feedback. In contrast, anxious individuals may exhibit less variability in error monitoring levels, rigidly deploying heightened error monitoring regardless of task/feedback instructions. Of note, this interpretation is also consistent with Attentional Control Theory (Eysenck et al., 2023), which poses that anxious individuals deploy heightened error monitoring levels as a compensatory strategy, in an effort to maintain performance and avoid errors—even when task instructions do not explicitly emphasize accuracy (Eysenck et al., 2023; Moser et al., 2013). If this hypothesis is correct, then even though studies commonly present both speed and accuracy feedback, it would specifically be the presence of speed feedback that drives the effect of block-level feedback on ERN-anxiety associations—as speed feedback would maximize error monitoring differences between anxious and non-anxious individuals. The current meta-analysis is unable to directly test this hypothesis, given that proportions of feedback type presented (speed vs. accuracy) are typically not computed nor reported. Moreover, as the feedback presented is typically contingent on participant performance (in line with the recommendations of Gehring et al., 2012), effects of feedback type and behavioral performance are fully confounded in most prior studies. Thus, to test whether the effect of block-level feedback on ERN-anxiety associations is driven specifically by speed feedback, a study is needed that manipulates not only whether feedback is presented, but also the type of feedback presented, in a within-subjects design.

#### No observed differences by stimulus type

We did not identify a significant effect of stimulus type on ERN-anxiety associations. However, we caution against drawing strong inferences based on this null effect. One reason for assessing possible effects of stimulus type on ERN-anxiety associations was because the CEMH theory of Moser and colleagues (2013) proposes that ERN-anxiety associations are driven by verbal rumination. Thus, if letter (vs. arrow) stimuli are more likely to prime verbal rumination, then one possibility is that ERN-anxiety associations would be stronger in studies employing a letter flanker task. However, it may be incorrect to assume that letter vs. arrow stimuli would differ substantially, if at all, in their ability to prime verbal rumination. Indeed, Lin and colleagues (2015) argue that even arrow stimuli, if horizontally arranged, may prime verbal rumination. A second reason for assessing possible effects of stimulus type on ERN-anxiety associations was that letter (vs.) arrow Flanker tasks differ in the relative strength of stimulus-response mappings, with stronger stimulus-response mappings for arrow stimuli due to a preexisting semantic-spatial compatibility (Kornblum et al., 1990). Thus, performing an arrow Flanker task might place less demands on working memory and allow for more efficient/automatic response preparation (Kornblum et al., 1990; Wilhelm & Oberauer, 2006), in turn impacting error monitoring and ERN-anxiety associations. Although no effect of stimulus type on ERN-anxiety associations was observed, we cannot rule out the possibility that orthogonal, yet opposing effects of letter and arrow stimuli on ERN-anxiety associations could have resulted in a null effect. Thus, further work is needed to carefully, and independently, manipulate Flanker stimuli along separate dimensions to provide a more definitive test of whether Flanker stimulus type influences ERN-anxiety associations. Nonetheless, at a practical level, and at least based on the current data, there is no evidence to suggest that the choice of letter vs. arrow stimuli will have a strong impact on observed ERN-anxiety associations.

### Limitations and future directions

The current study is not without limitations. Studies included in our meta-analysis were drawn from an existing database of studies (Saunders & Inzlicht, 2020), which allows for more direct comparison of how our moderation results compare with the results reported by Saunders and Inzlicht (2020). Moreover, we have no reason to believe that restricting our analyses to this set of studies compromises the results or conclusions drawn. Nonetheless, future work could improve on the current study by extending the analyses to include additional studies beyond those included in the database of studies published by Saunders and Inzlicht (2020). Relatedly, the current study investigated three task parameters of interest (stimulus type, response type, and block-level feedback), however, future work could expand on the set of parameters examined in the current report. The overarching limitation of all meta-regression analyses is that they must rely on previously collected data, and despite attempts to control for variation across studies and isolate the effects of moderating variables, there is inevitable variation across studies that is unaccounted for. Given meta-analytic evidence that response type and block-level feedback moderate ERN-anxiety associations, what is needed now are studies that directly manipulate these parameters at the within-study level, seeking to replicate the effects reported here. Relatedly, the goal of the current meta-analysis was not to provide a direct test of prior theory, but only to provide evidence, at the meta-analytic level, that specific task parameters can impact the magnitude of ERN-anxiety associations. Given that multiple theories can explain the moderating effects of response type and block-level feedback, future work could introduce further manipulations at the within-study level to disentangle the predictions of competing theories.

### Conclusions

The primary findings of this study are that two-handed (vs. one-handed) Flanker tasks, as well as those with (vs. without) block-level feedback, exhibit stronger ERN-anxiety associations. We did not find any evidence to suggest that stimulus type (arrow vs. letter) impacts ERN-anxiety associations among Flanker task studies—although we urge caution when interpreting this null result. At a practical level, the results suggest that researchers seeking to maximize ERN-anxiety associations should employ a two-handed arrow/letter Flanker task with block-level feedback. However, studies directly manipulating these task parameters (stimulus type, response type, and block-level feedback) are needed to confirm such recommendations. At a theoretical level, the results also lead to several testable hypotheses as to why response type and block-level feedback impact the magnitude of ERN-anxiety associations. Future work should test these hypotheses to advance theoretical understanding of ERN-anxiety associations. Finally, the results underscore the importance of studies carefully reporting task parameters to improve reproducibility, as variations in these parameters can help explain some of the inconsistencies found across the literature.

## Data Availability

Given that the initial list of studies and all effect sizes included in the current meta-analysis were sourced from an open data set of published/unpublished ERN-anxiety effect sizes created by Saunders and Inzlicht (2020), we refer readers to their original manuscript and corresponding OSF website (https://osf.io/r7dvc/). Additional data constructed as part of the current study (novel coding of task parameters), as well as all analysis code, can be found on the current paper’s companion GitHub website (https://github.com/NDCLab/flanker-ern-meta).

## Acknowledgments

We wholeheartedly thank Saunders and Inzlicht (2020) for their well-documented, carefully-crafted, and open database of published and unpublished ERN-anxiety effect sizes. Their publication of this open database and commitment to open science have made the current study possible.

## Supplementary materials

Additional analyses that go beyond the scope of the main analyses are provided in the supplementary materials.

