## Supplemental Material for "Flanker task parameters are related to the strength of association between the ERN and anxiety: a meta-analysis"

### Similar Meta-Regression Results When Excluding "Modified" Flanker Task Studies

As described in the main text, we coded studies in terms of whether they employed a Flanker task, as well as whether the task structure was "modified" to include additional stimuli or response demands that are not typical of most Flanker task studies. A subset of 5 studies were identified as being a "modified" version of the standard Flanker task structure. These modified Flanker task studies were included in all primary analyses reported in the main text; however, we additionally re-ran all random effects meta-regression models testing for moderation after removing the modified Flanker task studies and obtained similar results. The results of these additional analyses are presented in Table S1.

Table S1. Effect of task parameters on effect sizes associated with ERN-anxiety associations after removing "modified" Flanker tasks

| | **Q** | ***p*** | **effect** | **95% CI** | **K** | **N** |
|---|---|---|---|---|---|---|
| **Stimulus type** | 0.019 (1) | 0.891 | | | | |
| **Arrow** | | | -0.204 | -0.274, -0.135 | 29 | 2188 |
| **Letter** | | | -0.214 | -0.328, -0.099 | 12 | 835 |
| **Response type** | 5.404 (1) | 0.020* | | | | |
| **One-handed** | | | -0.109 | -0.204, -0.013 | 13 | 1128 |
| **Two-handed** | | | -0.248 | -0.316, -0.180 | 27 | 1836 |
| **Block-level feedback** | 13.402 (1) | <.001* | | | | |
| **Without** | | | -0.089 | -0.148, -0.029 | 13 | 1114 |
| **With** | | | -0.235 | -0.285, -0.184 | 23 | 1586 |

*Note: *denotes significance. ERN, error-related negativity.*

### Similar Meta-Regression Results When Analyzing Only Anxious Apprehension/Worry Studies

To maximize the number of included studies, for all primary analyses reported in the main text we included studies regardless of the anxiety measure used. However, we also performed an exploratory set of analyses in which we additionally re-ran all random effects meta-regression models testing for moderation when restricted to studies employing anxiety measures mapping onto the anxious apprehension/worry component of anxiety and obtained similar results. The results of these additional analyses are presented in Table S2.

Table S2. Effect of task parameters on effect sizes associated with ERN-anxiety associations for studies measuring only anxious apprehension/worry

| | Q | *p* | effect | 95% CI | K | N |
|---|---|---|---|---|---|---|
| **Stimulus type** | 0.044 (1) | 0.834 | | | | |
| **Arrow** | | | -0.229 | -0.289, -0.169 | 23 | 1720 |
| **Letter** | | | -0.241 | -0.342, -0.141 | 8 | 725 |
| **Response type** | 3.521 (1) | 0.061 | | | | |
| **One-handed** | | | -0.141 | -0.241, -0.041 | 6 | 657 |
| **Two-handed** | | | -0.251 | -0.308, -0.195 | 24 | 1729 |
| **Block-level feedback** | 9.495 (1) | .002* | | | | |
| **Without** | | | -0.101 | -0.179, -0.023 | 6 | 653 |
| **With** | | | -0.248 | -0.299, -0.196 | 21 | 1530 |

*Note: *denotes significance. ERN, error-related negativity.*

*Supplement to: "Flanker task parameters are related to the strength of association between the ERN and anxiety: a meta-analysis"*

*George A. Buzzell, Yanbin Niu, Emily Machado, Renata Dickinson, Jason S. Moser, Santiago Morales, Sonya V. Troller-Renfree*

**Evidence of Publication Bias Exclusively for Studies Without Block-Level Feedback**

Within the main text, we report the results of a random-effects meta-regression model, which found that block-level feedback significantly moderated ERN-anxiety associations ($p < .001$). For this model, the predicted effect size was larger (more negative) for studies with block-level feedback, -0.237 (95% CI: -0.285, -0.189), as opposed to studies without block-level feedback, -0.088 (95% CI: -0.147, -0.029). However, we also report in the main text that Egger's test of publication bias was significant ($p = 0.030$) for the studies employed to test the influence of block-level feedback. Here, we further assessed possible source(s) of publication bias (via Egger's test) in separate models fit at each level of the moderator (subgroup analysis for studies with vs. without block-level feedback). Additionally, we estimated the bias-corrected effect size for each subgroup via the trim-and-fill procedure (Duval & Tweedie, 2000). Results of these analyses are presented below:

Evidence for possible publication bias was only found within the subgroup of studies without block-level feedback, as indicated by a significant Egger's test exclusively for the subgroup without ($p = .014$) vs. with ($p = .759$) block-level feedback. Correcting for bias via the trim-and-fill procedure yielded a non-significant effect size estimate for the subgroup of studies without block-level feedback: -0.089 (95% CI: -0.197, 0.019). In contrast, the bias-corrected estimate for the subgroup of studies with block-level feedback remained significant: -0.220 (95% CI: -0.267, -0.174). Collectively, these additional analyses suggest that the possible presence of publication bias is exclusive to studies without block-level feedback, but does not qualitatively change the results and conclusions reported in the main text.

*Supplement to: "Flanker task parameters are related to the strength of association between the ERN and anxiety: a meta-analysis"*

*George A. Buzzell, Yanbin Niu, Emily Machado, Renata Dickinson, Jason S. Moser, Santiago Morales, Sonya V. Troller-Renfree*
